# Ligand-Receptor Interactions for Cell-Cell Communication Analysis in Rat, Chicken, Pig, and Monkey Single-Cell and Spatial Transcriptomics

**DOI:** 10.1101/2024.10.12.617999

**Authors:** Sheng Chen, Yan Peng, Xiaoling Zhang, Ting Jiang, Binbin Fang, Pengyue Zhang, Yanze Li, Yonghong Ren, Yimin Sun

**Affiliations:** Scientific Research Cooperation Center, CapitalBio Technology, Beijing 100176, China; National Engineering Research Center for Beijing Biochip Technology, Beijing 102206, China

## Abstract

Cell-cell communication is a frequently used analysis approach in single cell RNA and spatial transcriptomics, and many tools like CellPhoneDB, CellChat and stLearn have been developed. Ligand-receptor interactions are the core of cell-cell communication analysis. Since receptor-ligand and even protein-protein interactions were focus on humans and mice research, curated human and mouse receptor-ligand databases have been established, cell-cell interactions for these two species single-cell RNA sequencing data can be directly analyzed. However, for rats, chickens, pigs, monkeys, and other species, cell-cell interaction analysis is often implemented through orthologous gene mapping, due to the lack of curated ligand-receptor interaction databases for these species.

We collected cell-cell interaction data mainly from KEGG for rats, chickens, pigs, and monkeys, and extended the data from Reactome and IntAct. Then, by using CellChatV2 with our collected rat ligand-receptor interactions and CellChatV2’s own mouse data, respectively, we analyzed 10x Genomics rat public scRNA data, and found that 70 significantly ligand-receptor interactions from the mouse analysis result were also significantly in rats. We also obtained some chicken, pig, and monkey scRNA data from published literature, and analyzed cell-cell interactions using our collected ligand-receptor interactions for these species, and it was proved that our data is reliable and useful. Lastly, we have transformed the ligand-receptor interactions for rat, chicken, pig, and monkey species into the CellChatDB format, which enables swift and straightforward analysis of cell-cell communication in single-cell and spatial data of these four species.

All the ligand-receptor interaction datasets for rats, chickens, pigs, and monkeys, as well as the program codes, are available at https://github.com/qingchen36/ligand-receptor. Using our program, one can rapidly obtain receptor-ligand interaction data for other species.

## Introduction

Cell-cell communication mediated by ligand-receptor interactions is crucial for coordinating diverse biological processes, such as development, differentiation, and responses to infection. Single-cell RNA sequencing and spatial transcriptomics have been powerful tools for investigating cellular biology at an unprecedented resolution, enabling the characterization of cellular heterogeneity. After major or sub cell-type annotation, cell-cell communication has been one of the most popular analysis modules in scRNA and spatial transcriptomics research[1]. There have been many cell-cell communication analysis methods for scRNA, such as iTALK[2], CellTalker[3], NicheNet[4], CellPhoneDB[5], and CellChat[6], and for spatial transcriptomics, such as stLearn[7], SpatialDM[8], spaCI[9], and CellChat V2[10]. These cell-cell communication methods or tools were mainly used with human data, but there is no tool that can directly analyze rat, chicken, pig, and monkey single-cell and spatial transcriptomics data.

CellPhoneDB only had human-sourced ligand-receptor pairs, and CellChat also had mouse and zebrafish data, but for rat, chicken, pig, and other species, there are no pre-existing ligand-receptor interaction datasets. For these species without ligand-receptor interaction datasets, before cell-cell communication analysis, researchers usually map the gene names to their nearest species by orthology, just as Jiao Qu published “A reference single-cell regulomic and transcriptomic map of cynomolgus monkeys” in Nature Communications in 2022 mentioned, converting monkey genes to human genes[23]. Due to the incompleteness of homologous gene databases, it was inevitable that some ligand and receptor genes could not be orthologized to human or mouse, especially when studying species like chickens or pigs. Most cell-cell communication inference can be conceptualized as a form of co-expression analysis, where the expression pattern of a ligand from one cell type and a paired receptor from another cell type is considered. Thus, ligand-receptor interaction datasets are the fundamental core of cell communication analysis, and collecting and organizing ligand-receptor interaction data is highly beneficial for scRNA and spatial transcriptomics research in these species.

CellphoneDB is a publicly available repository of human-curated receptors, ligands, and their interactions, and its ligand-receptor interactions were also the main source data for other cell-cell communication tools, such as CellChat V1. Studying the construction and development process of CellphoneDB can facilitate the establishment of databases for species like rats and chickens. CellphoneDB V1 data was first published in Nature in 2018[5], and it has been updated to V5 now. CellphoneDB defines ligands as secreted proteins, mainly from cytokines, hormones, growth factors, and immune-related proteins, and receptors as those that must be transmembrane. It also explicitly states that collecting the composition of receptor/ligand complexes and the interactions between ligand-receptor complexes are more in line with the actual circumstances of cell communication[11]. CellphoneDB V3 added ligand-receptor interactions from the Wnt signaling pathway[12], and V4 added non-protein receptor interactions, which are not used by default in scRNA cell-cell communication analysis[13]. CellphoneDB ligand-receptor interactions were mainly curated from high-confidence protein-protein interaction databases and published literatures.

If we want to establish a database containing rat, chicken, pig, and monkey ligand-receptor interactions by following the approach of CellphoneDB, firstly, more protein-protein interactions need to be researched, and then ligand-receptor pairs can be selected from them. The IntAct[14] database is a high-confidence protein-protein interaction database that collects evidence for molecular interactions and maintains the resource from iMEX databases, such as MatrixDB[15], MINT[16], and I2D[17]. In IntAct, protein-protein interactions covered about 2.6k rat proteins, which is a significant decline compared to the7.7k mouse proteins and 18k human proteins. For chicken, pig, and monkey, there are almost no collected protein-protein interactions, so selecting ligand-receptor interactions from existing protein-protein interaction databases is difficult to implement.

CellChat v2 included the direct incorporation of spatial locations of cells and can also infer spatially proximal cell-cell communication[10]. CellChat v2 expanded on more than 1000 protein and non-protein interactions based on peer-reviewed literature and existing databases such as CellPhoneDB[5] and NeuronChatDB[18]. Both CellChat and CellPhoneDB classify ligand-receptor interaction pairs into four categories: Cell-Cell Contact, ECM-Receptor, Secreted Signaling, and Non-Protein Signaling. Since non-protein Signaling ligand-receptor interactions were not used in scRNA and spatially resolved transcriptomics cell–cell communication analysis, we counted the other three categories of ligand-receptor data in CellChat and found that there were 1507 of 2378 mouse ligand-receptor pairs from KEGG Pathway[19], and 1416 of 2241 human ligand-receptor pairs from KEGG Pathway[19].

KEGG PATHWAY is a collection of manually drawn pathway maps representing our knowledge of molecular interactions, reactions, and relation networks[19]. It has always been a pivotal database for functional enrichment analysis in rat, chicken, pig, monkey, and other species. Therefore, through the curation of KEGG Pathways, it becomes feasible to collect a considerable number of ligand-receptor interactions for rat, chicken, pig, and monkey.

## Methods

### ligand-receptor interactions collection

We only curated ligand-receptor interaction pairs from 11 selected KEGG Pathways (details in Table 1) by downloading the KGML files for each species’ pathway through the KEGG API, and our data processing adhered to the following rules:

**Table 1.**
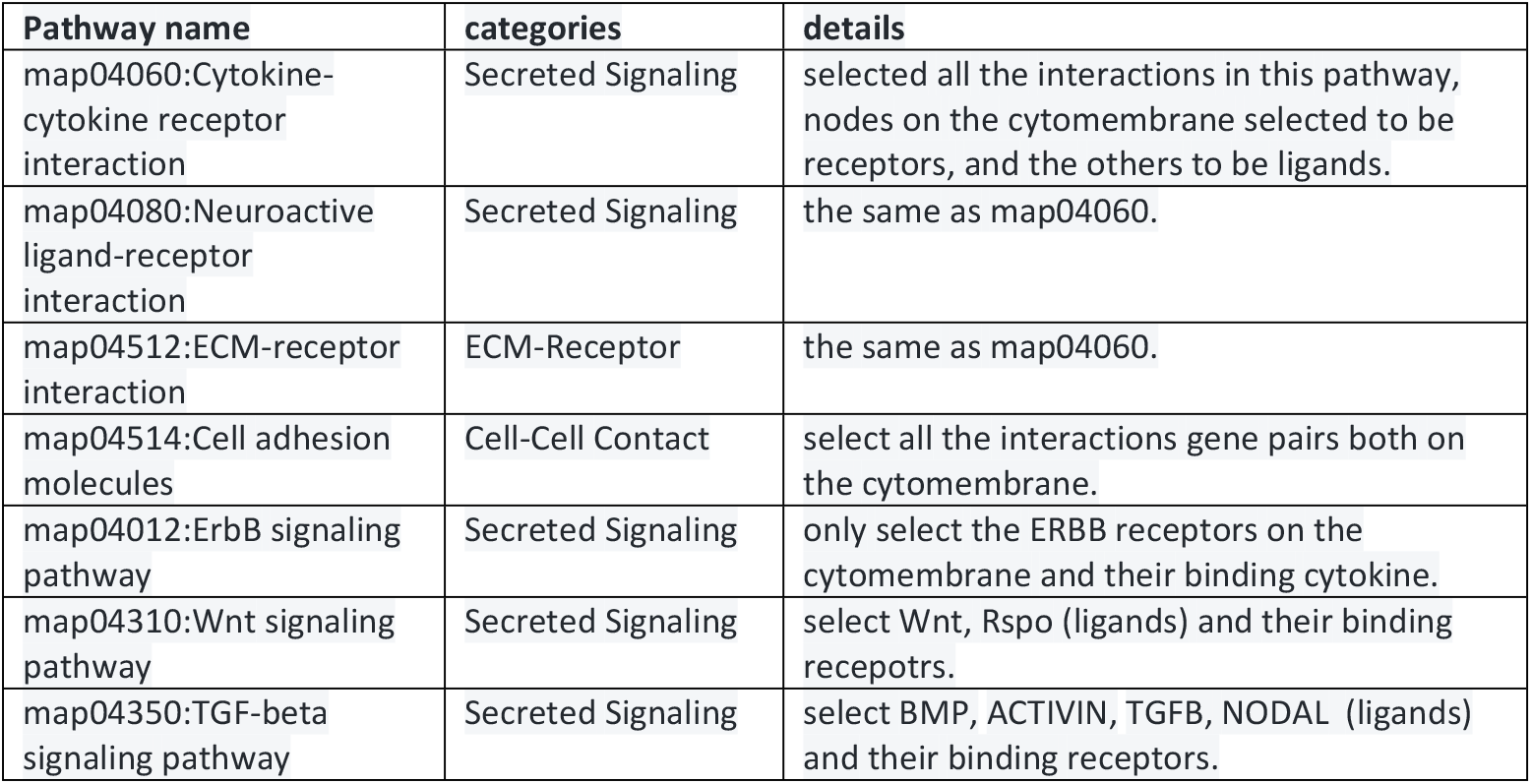
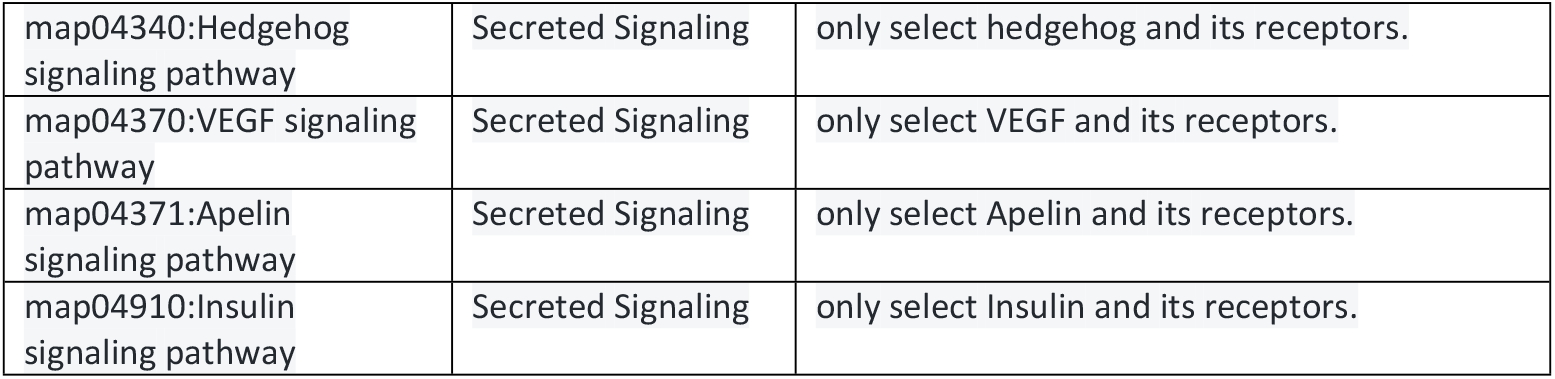
11 KEGG Pathways for ligands-receptors pairs collection.

1. Except for map04514 (cell adhesion molecules), for the other 10 pathways, we only select receptors located on the cytomembrane, and their binding ligand genes must be extracellular.
2. In the KGML files, if a node type is equal to “gene”, it indicates a sole node, ligand nodes and receptor nodes can be distinguished based on the relation direction. Each gene in ligand and receptor nodes was respectively selected to construct pairwise ligand-gene and receptor-gene interactions, when a ligand node or a receptor node annotated by more than one gene.
3. In the KGML files, if a node type is equal to “group”, it indicates a complex node. Each gene in each subunit node was respectively selected to construct all possible forms of gene complexes. Then, following rule b, we obtain ligand-gene and receptor-complex interactions, ligand-complex and receptor-gene interactions, or ligand-complex and receptor-complex interactions.
4. We download protein-protein interactions from IntAct[14] and Reactome[20], compile lists of ligand genes and receptor genes based on the 11 KEGG pathways, and then expand the ligand-receptor pairs from IntAct and Reactome.

### public single cell RNA data

The rat single-cell RNA data was downloaded from the 10x Genomics public datasets (https://www.10xgenomics.com/datasets/10-k-rat-pbm-cs-multiplexed-2-cm-os-3-1-standard-6-0-0). The chicken skeletal muscle single-cell RNA data [21] was downloaded from the Genome Sequence Archive in BIG Data Center, under accession number CRA002353 (https://bigd.big.ac.cn/gsa/). The pig skin cell scRNA-seq data [22] was downloaded from the GEO database, under accession code GSE166561. The monkey kidney scRNA-seq data [23] was downloaded from the URL https://doi.org/10.5281/zenodo.5881495.

### cell–cell communication analysis with CellChat

We transformed our collected ligand-receptor interactions for rat, chicken, pig, and monkey into the CellChat format, and then used CellChat V2 to analyze cell–cell communication for each of the rat, chicken, pig, and monkey scRNA datasets.

There were 26,309 protein coding genes in the mouse NCBI GeneInfo (version 20240521), while the rat NCBI GeneInfo (version 20240521) contained 23,280 protein coding genes, approximately 77.5% (18,048/23,280) of the gene symbols were consistent between mouse and rat. Therefore, for the rat data, we also ran CellChat using the CellChatV2 own mouse ligand-receptor interactions dataset. The usability of our curated ligand-receptor interactions dataset was assessed by comparing the common significant ligand-receptor pairs for every source-target cluster pair between rat and mouse.

## Results

### ligand-receptor interactions for chicken, monkey, pig and rat

The statistics of categories and sources for rat, monkey, chicken, and pig ligand-receptor interaction annotations can be seen in Fig. 1, above all, we collected 2,564 rat, 2,462 monkey, 1,968 chicken, and 2,620 pig ligand-receptor interaction pairs, which were mainly sourced from the KEGG Pathway database, and there were no monkey or pig ligand-receptor interactions, and only 12 rat and 3 chicken ligand-receptor interactions sourced from IntAct(more details in Supplementary Table 1). This also demonstrates that building a receptor-ligand interaction database for these four species from high-quality protein-protein interaction databases using the CellPhoneDB approach is virtually impossible.

**Fig 1.**
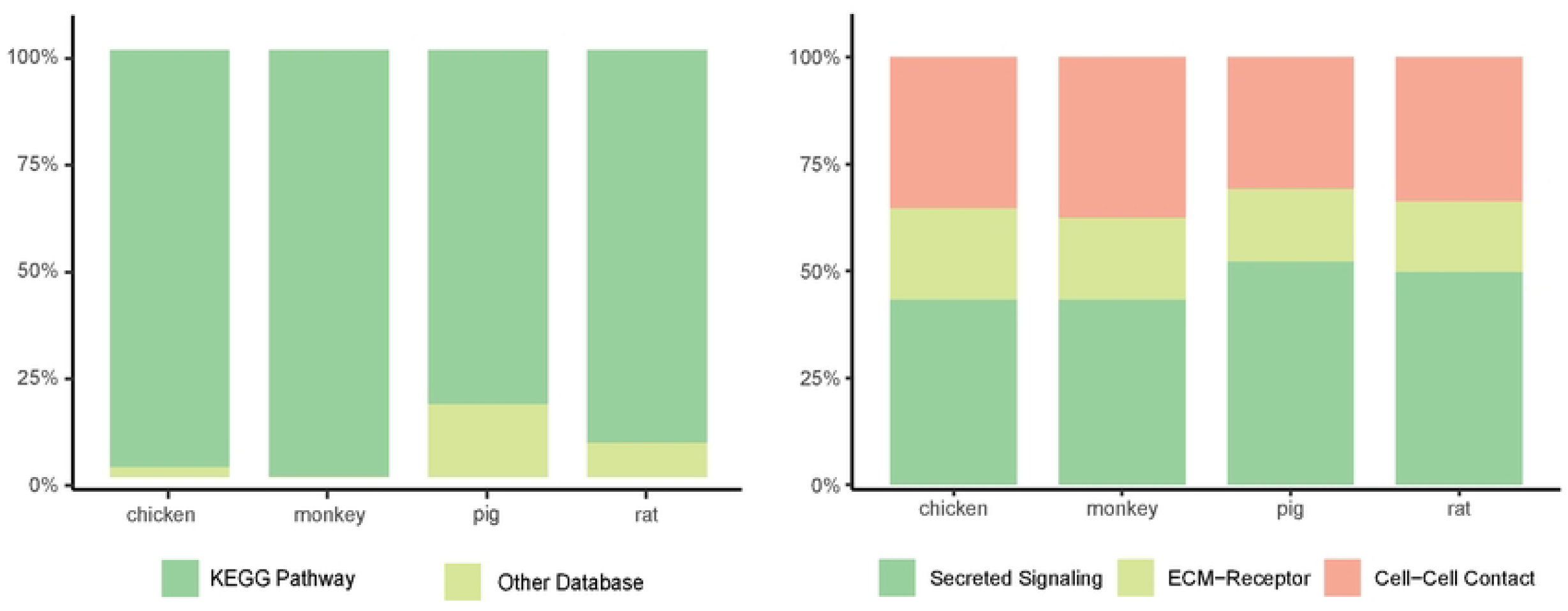
ligands-receptors pairs category and source database statistic

Additionally, the CellChat-format text and Rdata files containing the monkey, chicken, pig, and rat ligand-receptor interaction data are available at https://github.com/qingchen36/ligand-receptor/tree/main/CellChatDB. As a hint, the content of cofactor.xls files under each species directory were empty, as the existence of these files are essential for building the CellChat database.

### rat cell–cell interactions analysis

Non-protein receptor interactions were not used in the scRNA cell–cell communication analysis, and approximately 77.5% of rat gene symbols were the same as mouse. 92% of our collected rat ligand-receptor interactions were from KEGG Pathway. For the 10x Genomics public rat scRNA data, 5,380 cells were classified into 10 clusters. Firstly, we retained the KEGG-sourced receptor-ligand interactions from the CellChatV2 mouse databases and analyzed rat cell–cell interactions using the retained mouse databases with the CellChatV2 software. As shown in Fig. 2, we obtained a total of 70 significantly correlated ligand-receptor interactions between different cell clusters and within them. The details can be found in Supplementary Table 2. Secondly, we analyzed rat cell–cell interactions using our collected rat databases with the CellChatV2 software, and we obtained 348 significantly correlated ligand-receptor interactions between different cell clusters and within them(Fig. 3), the details can be found in Supplementary Table 3. Finally, we compared the results obtained from these two methods and found that all 70 ligand-receptor interactions from the mouse were also significant in the rat. This means that for each of these 70 interactions, their source cluster, target cluster, ligand, and receptor were the same in both rat and mouse results, the details can be found in Supplementary Table 4.

**Fig 2.**
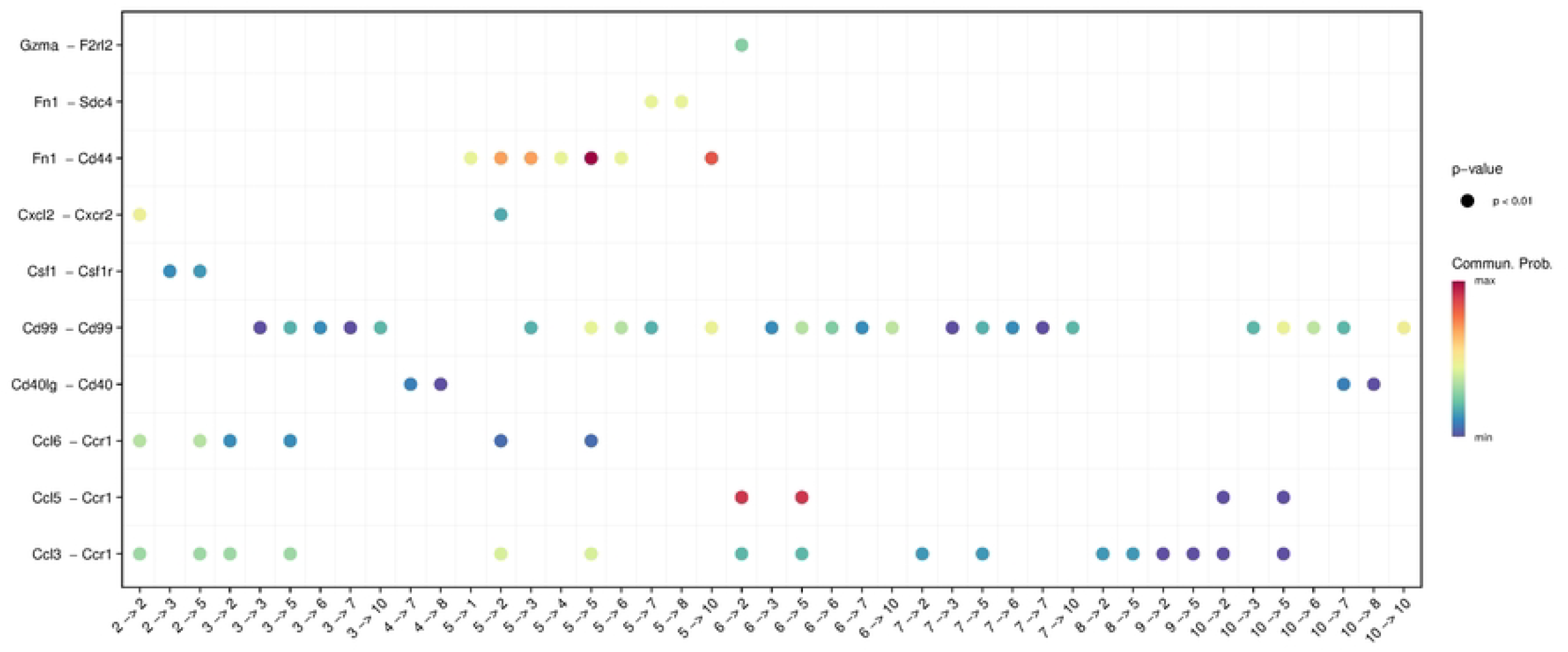
Rat cell–cell interactions analysis using mouse ligand-receptor interactions from CellChat, the abscissa in the figure represents the cell cluster from source to target, and the ordinate represents the ligand-receptor interaction pairs.

**Fig 3.**
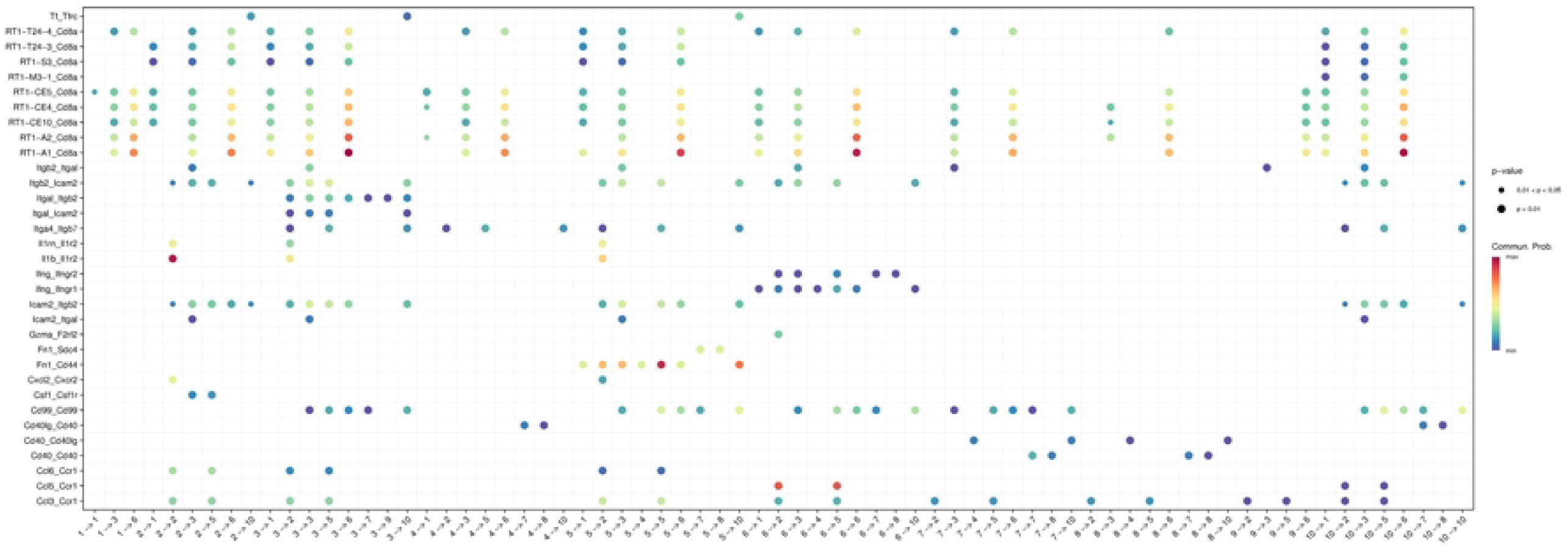
Rat cell–cell interactions analysis using our collected rat ligand-receptor interactions, the abscissa in the figure represents the cell cluster from source to target, and the ordinate represents the ligand-receptor interaction pairs.

### monkey cell–cell interactions analysis

Jiao Qu and Dijun Chen built “A reference single-cell regulomic and transcriptomic map of cynomolgus monkeys” in 2022. They collected single-cell RNA data from 16 monkey organs, annotated the cell types, and analyzed cell–cell interactions using CellPhoneDB by converting monkey genes to their corresponding human homologous genes. Since the keyword of we collected receptor-ligand interactions data was gene symbol, we downloaded monkey scRNA data, after extracting the kidney single-cell data, we converted the ensemble gene IDs in the expression matrix to their corresponding gene symbols. Then, we analyzed cell–cell interactions using our own collected data with CellChatV2 and obtained 111 significantly correlated ligand-receptor interactions(Fig. 4). The details can be found in Supplementary Table 5.

**Fig 4.**
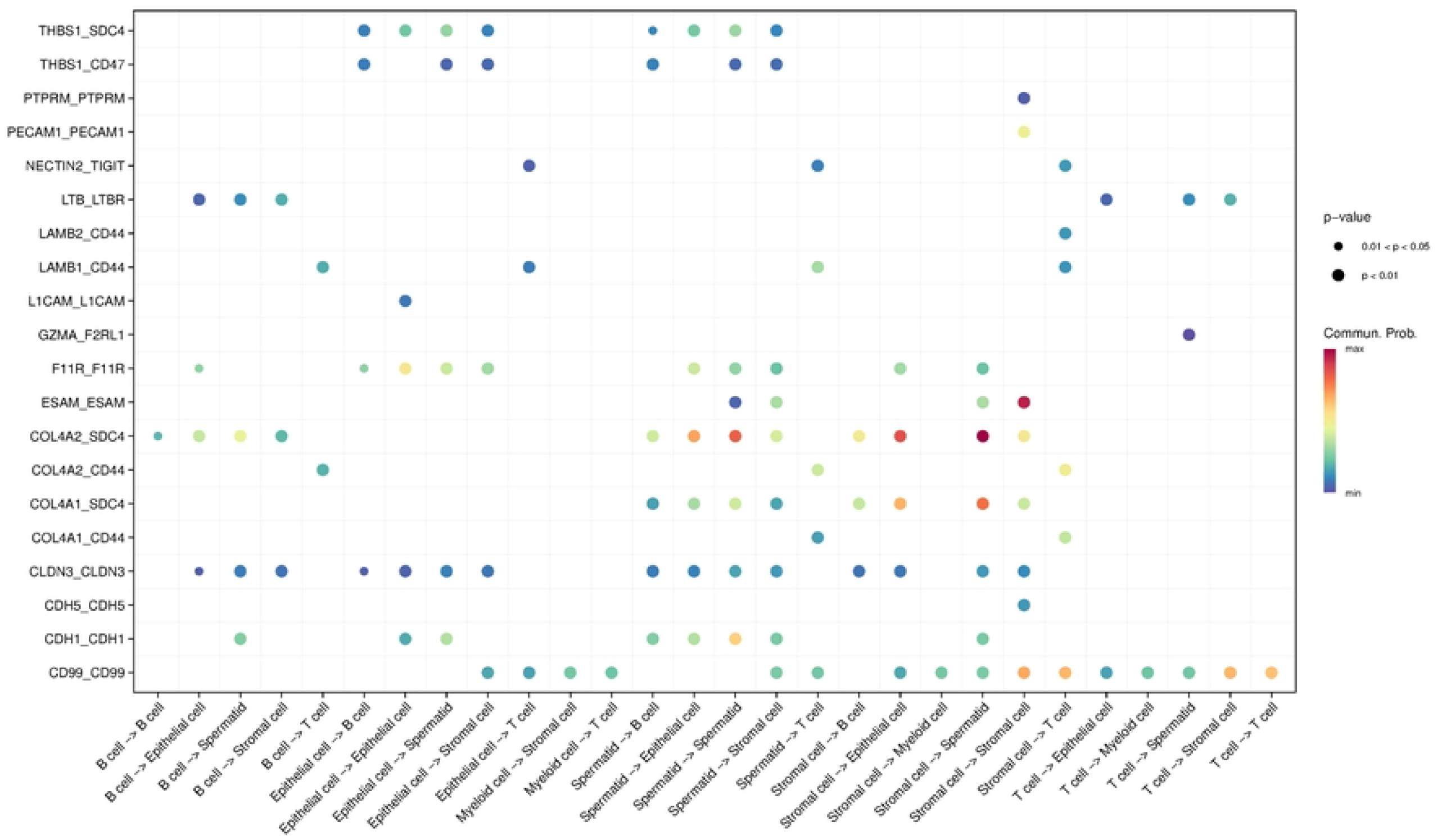
monkey cell–cell interactions analysis using our collected monkey ligand-receptor interactions databases, the abscissa in the figure represents the cell type from source to target, and the ordinate represents the ligand-receptor interaction pairs.

### chicken and pig cell–cell interactions analysis

Due to the limitations of homologous databases, even for two closely related species, there would be some genes that cannot be successfully converted. For species like chicken and pig, which are more distantly related to human or mouse, using homologous genes would lead to more information loss. Moreover, homologous databases are also predictive data, so the credibility of cell interaction analyses based on homologous genes would also be compromised. Therefore, using the methodology presented in this paper, curated chicken and pig ligand-receptor interactions from KEGG Pathway for cell interaction analysis is a more suitable approach. The cell-cell interactions analysis result between the chicken skeletal muscle related eight cell types defined by Li J [21]is shown in Fig. 5. and the cell-cell interactions analysis result between the pig skin related ten cell types defined by Han L [22]is shown in Fig. 6. In conclusion, our method provides a comprehensive analysis of cell interactions in these species.

**Fig 5.**
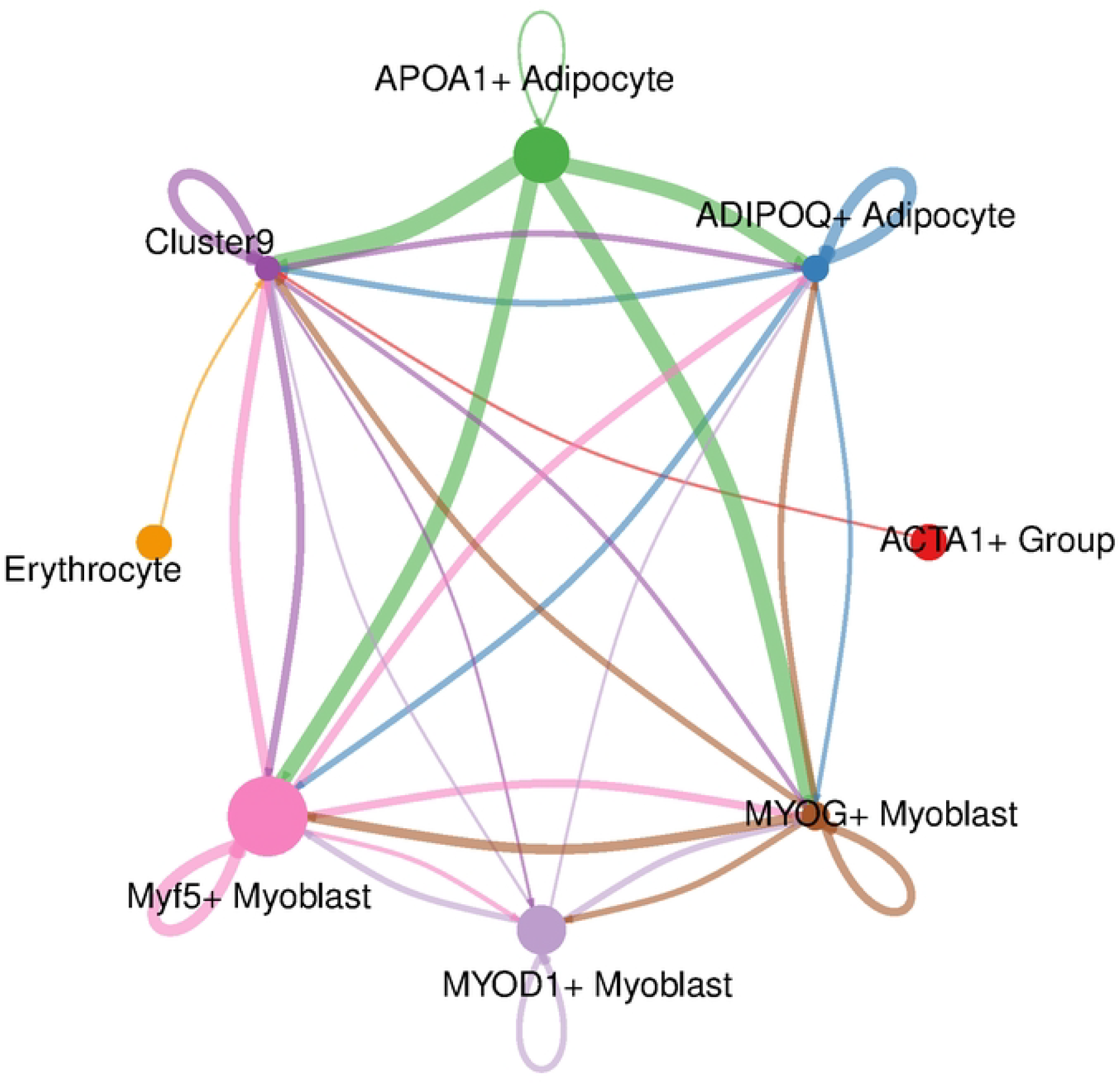
chicken cell–cell interactions analysis, the node in the network represents the cell type, the edge represents the interaction between two cell types, and the width of the edge represents the number of ligand-receptor pairs.

**Fig 6.**
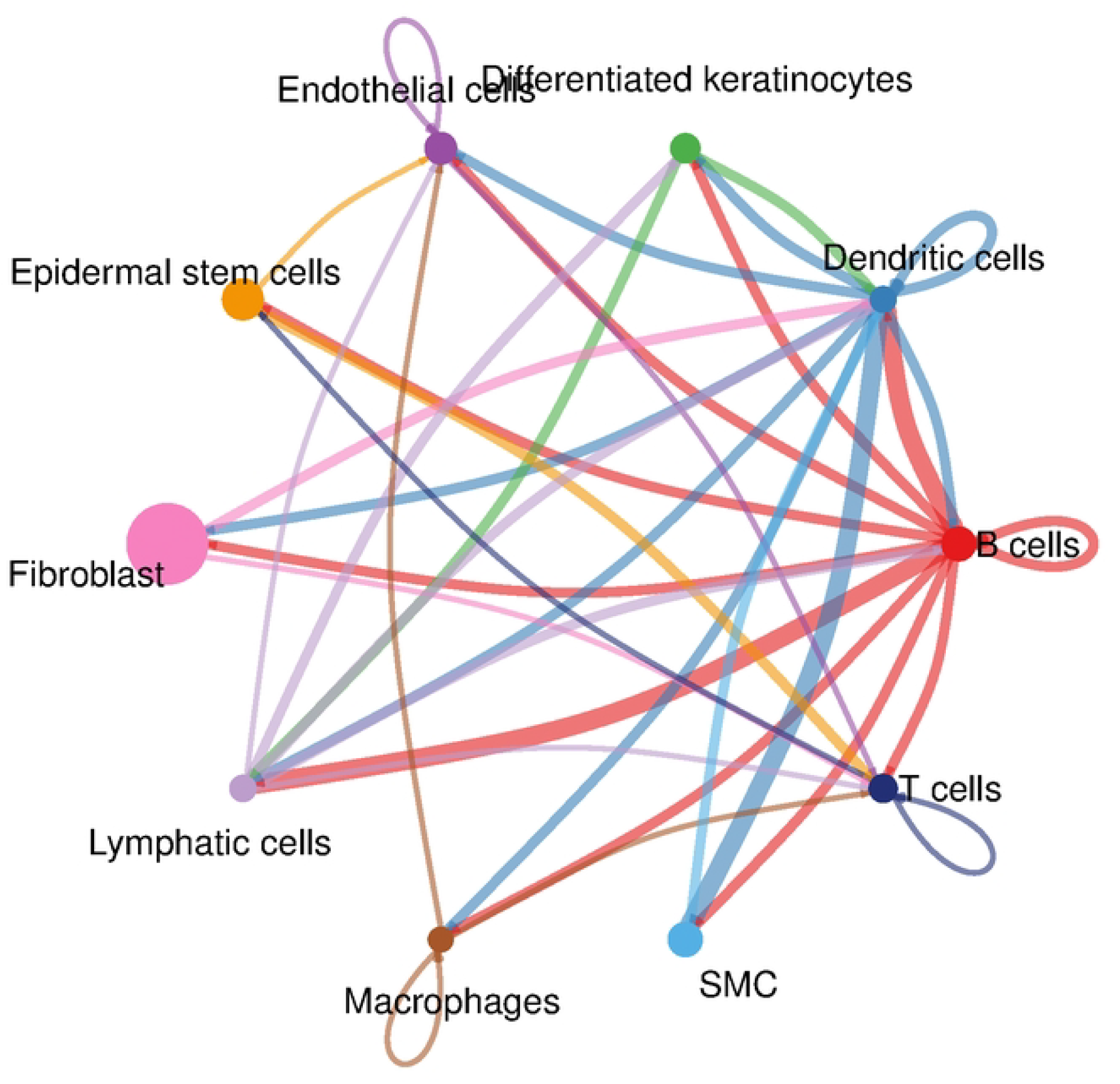
pig cell–cell interactions analysis, the node in the network represents the cell type, the edge represents the interaction between two cell types, and the width of the edge represents the number of ligand-receptor pairs.

All the curated data and the script files used in our analyses are available on https://github.com/qingchen36/ligand-receptor. Furthermore, our method can also be extended to collect receptor-ligand interactions data for other species.

## Discussion

For the rat and monkey species, previously, due to the absence of a high-confidence, well-curated database of receptor-ligand interactions, before cell-cell interaction analysis, we had to map rat genes to their nearest mouse genes and monkey genes to human genes. In this paper, we curated KEGG Pathway data, enabling us to directly analyze cell-cell interactions in rats and monkeys without the need for orthologous mapping. Furthermore, by comparing the rat single-cell data with two results: using our rat database and CellChat’s mouse database, we found that all the cell-cell interaction pairs identified based on the mouse receptor-ligand database were also detected in the results from our rat database. This indicates that our curated rat receptor-ligand interaction data is valid and reliable. Additionally, utilizing our rat database yields more receptor-ligand interaction pairs, providing more information for downstream experiments.

Homologous databases always were predictive data, so the credibility of cell interaction analyses based on homologous genes would also be compromised. For species like chicken and pig, which are more distantly related to human or mouse, using homologous genes would lead to more information loss. In this paper, we provide a method for rat, monkey, chicken and pig ligand-receptor interactions collection, which has been successfully applied to the cell-cell interaction analysis in these four species single cell RNA data, demonstrating the universality of this method. This method can also be extended to ligand-receptor interactions collection for other species.

Compared to the human and mouse ligand-receptor interaction databases, there is still a significant gap in the data for rats, monkeys, pigs, and chickens. Specifically, there is a lack of information about cofactors, soluble agonists, antagonists, co-stimulatory, and co-inhibitory membrane-bound receptors, which has been collected through experimental verification or reported in literature. Due to the absence of highly reliable cofactor information for these four species, we could consider supplementing this information using homology-based methods. However, this might introduce uncontrollable errors. Therefore, we have decided to only collect and organize the most core ligand-receptor interaction data.

## Code Availability

All the curated data and the script files used in our analyses are available on https://github.com/qingchen36/ligand-receptor.

## Acknowledgements

We extend our gratitude to the colleagues from the Bioinformatics Department for their assistance in this work, and I personally thank my family for their unwavering support of my professional endeavors.

## Author contributions

Project conception, Yan Peng, Ting Jiang, Sheng Chen, and Yanze Li; investigation, XL Zhang, Sheng Chen and Ting Jiang; data curation and analysis, Sheng Chen, Peng Yan; writing-review and editing, Sheng Chen, Peng Yan and YZ Li; funding acquisition, Yonghong Ren and Yanze Li. All authors read and approved the final manuscript.

